# CRISPR-CLEAR: Nucleotide-Resolution Mapping of Regulatory Elements via Allelic Readout of Tiled Base Editing

**DOI:** 10.1101/2024.09.09.612085

**Authors:** Basheer Becerra, Sandra Wittibschlager, Zain M. Patel, Ana P. Kutschat, Justin Delano, Eric Che, Anzhelika Karjalainen, Ting Wu, Marlena Starrs, Martin Jankowiak, Daniel E. Bauer, Davide Seruggia, Luca Pinello

**Affiliations:** Bioinformatics and Integrative Genomics PhD Program, Harvard Medical School, Boston, MA, USA; Broad Institute of MIT and Harvard, Cambridge, MA, USA; Molecular Pathology Unit, Krantz Family Center for Cancer Research, Massachusetts General Hospital Research Institute, Boston, MA, USA; Department of Pathology, Harvard Medical School, Boston, MA, USA; Division of Hematology/Oncology, Boston Children’s Hospital, Department of Pediatric Oncology, Dana–Farber Cancer Institute, Harvard Stem Cell Institute, Department of Pediatrics, Harvard Medical School, Boston, MA, USA; St. Anna Children’s Cancer Research Institute (CCRI), Vienna, Austria; CeMM Research Center for Molecular Medicine of the Austrian Academy of Sciences, Vienna, Austria

## Abstract

CRISPR tiling screens have advanced the identification and characterization of regulatory sequences but are limited by low resolution arising from the indirect readout of editing via guide RNA sequencing. This study introduces *CRISPR-CLEAR*, an end-to-end experimental assay and computational pipeline, which leverages targeted sequencing of CRISPR-introduced alleles at the endogenous target locus following dense base-editing mutagenesis. This approach enables the dissection of regulatory elements at nucleotide resolution, facilitating a direct assessment of genotype-phenotype effects.

## MAIN

While chromatin profiling, GWAS, and eQTL analyses propelled significant breakthroughs in identifying potential functional non-coding regions and variants in disease^1^, these methods do not provide mechanistic understanding of regulatory elements at the nucleotide level. In this context, CRISPR-based methods facilitate *in situ* perturbations of regulatory sequences, presenting opportunities to explore regulatory functions across various developmental stages and in numerous disease states in their native chromatin context ^2–6^. Current CRISPR screens commonly perform indirect readout of perturbations via guide RNA sequencing^7^. However, this readout is largely confounded by the editing efficiency of each guide RNA, and the enrichment of specific editing outcomes cannot be resolved. To address these limitations, we introduce *CRISPR-CLEAR* (Conveniently Linking Enriched Alleles to Regulation), a novel experimental and computational framework that enables precise determination of genotype-phenotype relationships at the nucleotide level, utilizing information from direct sequencing of alleles produced by base editing at regulatory regions (**Fig. 1a**).

**Figure 1:**
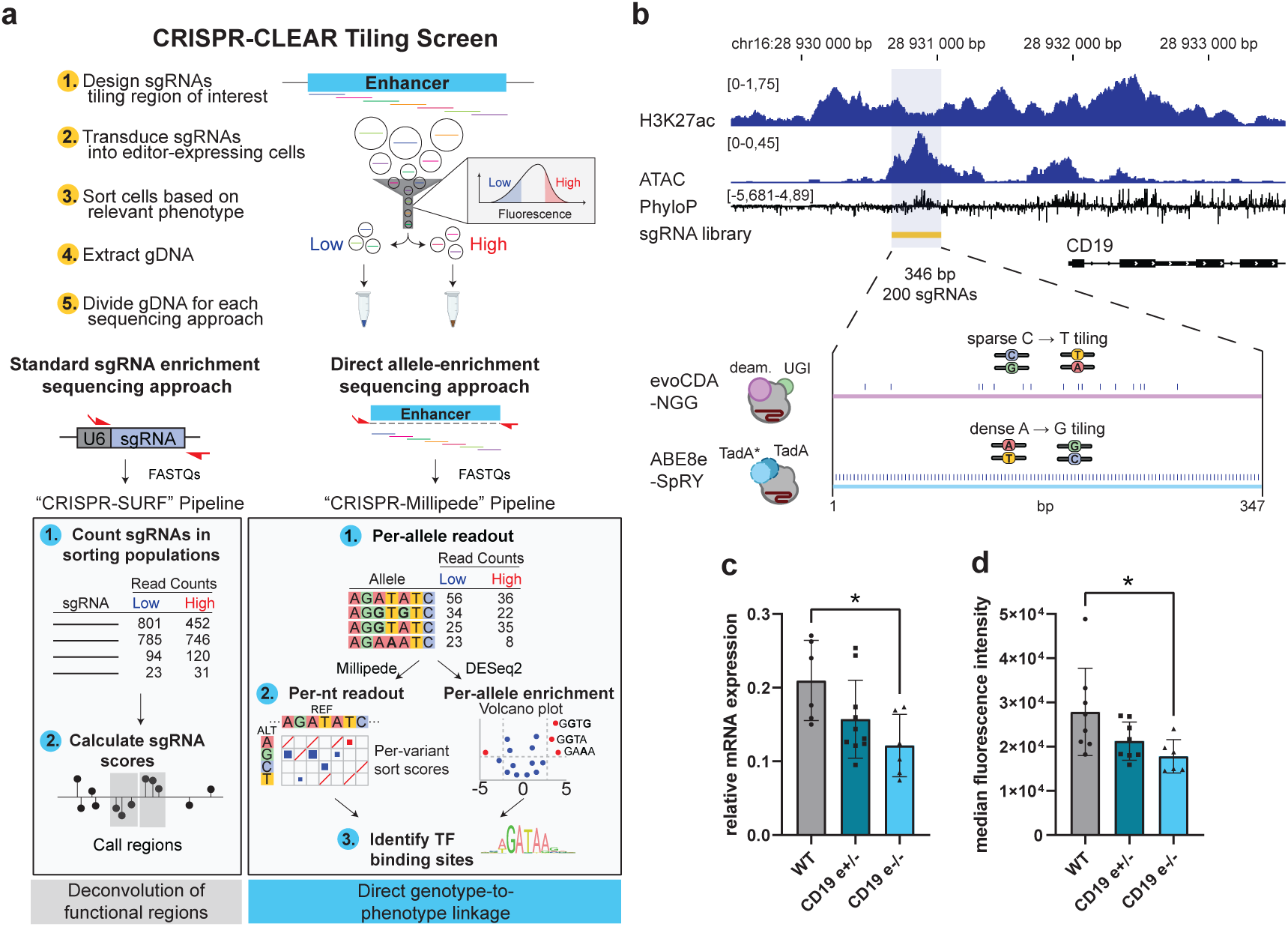
A base-editor tiling screen with allele-based readout. **a.** Comparison of *CRISPR-CLEAR* workflow with standard sgRNA enrichment sequencing approach. The workflow illustrates the key steps from guide RNA design to data analysis. First, cells stably expressing a base editor are transduced with a library of guide RNAs tiling the regulatory sequence. After editing, cells are FACS-sorted based on the expression of the target protein. Genomic DNA is extracted from sorted cells. Next-generation libraries are prepared to quantify sgRNA counts and to measure the distribution of edits at the endogenous sequence in the sorted population of cells. The left pathway shows the standard approach using sgRNA count-based readout and the *CRISPR-SURF* pipeline for deconvolution of functional regions. The right pathway depicts the *CRISPR-CLEAR* approach using direct allele-based readout and the *CRISPR-Millipede* pipeline, enabling precise genotype-to-phenotype linkage through per-allele and per-nucleotide analysis. **b**. A putative proximal enhancer is located upstream of the *CD19* promoter, based on H3K27ac and ATAC-seq of NALM6 cells, and sequence conservation. **c**. NALM6 clones with mono-or biallelic deletion of the *CD19* enhancer show reduction in CD19 mRNA levels. **d**. Wild-type and enhancer KO clones were stained with a CD19 antibody. Flow cytometry indicated a reduction in CD19 protein levels. One-way ANOVA, replicates are shown in circles/squares/triangles (n=6-10), * = p<0.05).

To demonstrate the efficacy of the *CRISPR-CLEAR* framework, we investigated a putative regulatory element upstream of the *CD19* gene, a B-cell marker and CAR-directed therapy target in leukemia. The candidate regulatory element was identified based on the presence of a highly-conserved, open chromatin region in proximity to the *CD19* gene (**Fig. 1b**), as indicated by ATAC-seq and H3K27ac ChIP-seq signal^8^. Deletion of the 346bp element resulted in reduced levels of CD19 mRNA (**Fig. 1c**) and protein (**Fig. 1d**). We designed a library of 200 guide RNAs tiling the *CD19* enhancer and performed screens using both cytosine (evoCDA^9^) and adenine (ABE8e-SpRY^10^) base editors in NALM-6 cells, a B-cell leukemia cell line. The ABE8e-SpRY base editor operates without a PAM requirement, therefore installing edits at high density at the *CD19* enhancer (**Supplementary Fig. 1a**) in contrast to the evoCDA base editor which is restricted by the –NGG proximal sequences (**Supplementary Fig. 1b**). While ABE8e performs A-to-G substitutions (**Supplementary Fig. 1c**), evoCDA is a spurious editor primarily introducing C-to-T substitutions along with different types of C-to-N substitutions and short *indels*, thereby increasing allele diversity (**Supplementary Fig. 1d)**.

After transducing the guides at low MOI, we cultured the cells for 7 days to allow for efficient editing and phenotype development. We then stained cells with a fluorescently conjugated antibody and performed fluorescence-activated cell sorting (FACS) in triplicates to sort cells into “CD19 positive” or “CD19 negative” populations. We targeted the *AAVS1* locus as negative control and the *CD19* exon 2 splice site as positive control (**Supplementary Fig. 1e**). For both the CD19 positive and CD19 negative sorted samples, we split the extracted genomic DNA for both targeted sequencing of the guide RNA cassette and the endogenously targeted region. This approach facilitated a direct and controlled comparative analysis between the standard guide count-based CRISPR screens and our proposed direct allele readout method, *CRISPR-CLEAR*.

To establish a baseline for comparison and evaluate the efficacy of the standard guide count-based CRISPR screens in identifying hotspot regions within the *CD19* regulatory element, we applied *CRISPR-SURF*^11^, a method specifically designed to analyzing such screens, to the guide RNA count data derived from both ABE8e and evoCDA base editors. Due to the unrestrictive PAM of ABE8e-SpRY, we have developed a guide RNA mapping method to handle “self-editing” at the guide RNA cassette and increase guide count recovery and power (**Supplementary Fig. 2a**). After calculating the enrichment of guide RNA read counts between the CD19 positive and CD19 negative populations, *CRISPR-SURF* identified one significant hotspot region at amplicon position 220-240 that was shared between evoCDA and ABE8e screenings, along with a second hit unique to the evoCDA screen at amplicon position 140-160 (**Fig. 2a**).

**Figure 2:**
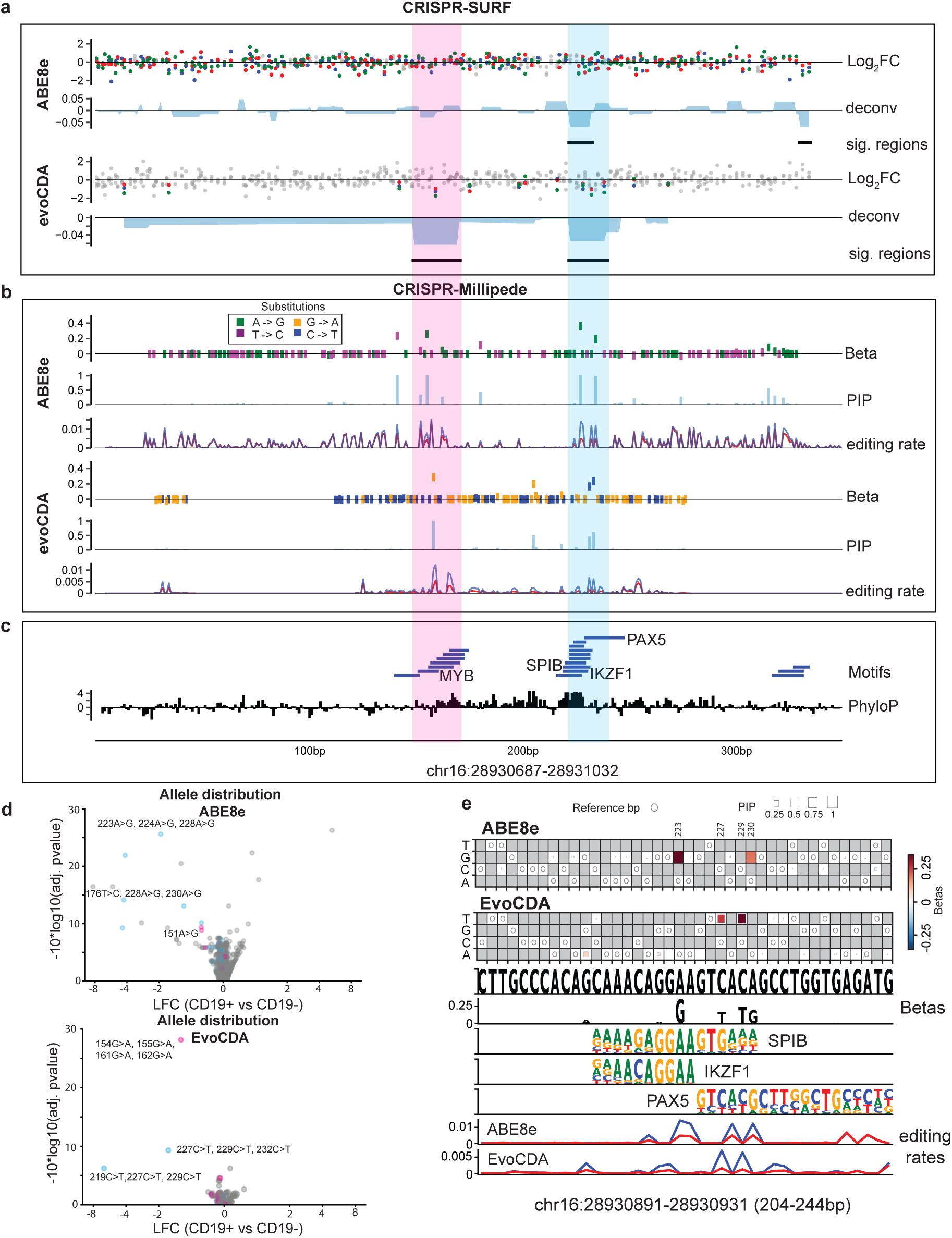
Direct measurement of regulatory potential using allele-based readout. **a.** *CRISPR-SURF* analyses. Top: plot showing the fold enrichment of sgRNA in CD19 positive versus CD19 negative cells in the ABE8e-SpRY screen. Scores of guides from each of three replicates are shown in red, blue, or green. Scores of negative control sgRNAs (20mers not containing editable adenines) are shown in grey. Deconvolution score track with the region called significant at positions 220-230 by *CRISPR-SURF* demarcated with a black segment below. Bottom: plot showing the fold enrichment of guides from the evoCDA screen. Negative control guides (20mers lacking a cytosine within the expected editing window, with a non-NGG PAM) are shown in grey. Deconvolution score track with the regions called as significant at positions 140-150 and 220-230 by *CRISPR-SURF* demarcated with a black segment below. **b.** *CRISPR-Millipede* analyses. From top to bottom: plot showing the effect sizes obtained for A>G (green) and T>C (purple) substitutions (covariates) at given positions in the sequence from the *CRISPR-Millipede* analysis of the alleles in the CD19 positive and CD19 negative sorted populations from the ABE8e-SpRY screen. Positive effect size indicates variants leading to lower CD19 expression. Track showing the posterior inclusion probabilities (PIP) for each of the covariates in the ABE8e-SpRY screen. Track showing the editing rate of A>G and T>C substitutions in the CD19 positive (red) and CD19 negative sorted populations (blue). Plot showing the effect sizes obtained for G>A (yellow) and C>T (blue) substitutions (covariates) at given positions in the sequence from the *CRISPR-Millipede* analysis of the alleles in the CD19 positive and CD19 negative sorted populations from the evoCDA screen. Track showing the posterior inclusion probabilities (PIP) for each of the covariate in the evoCDA screen. Track showing the editing rate of G>A and C>T substitutions in the CD19 positive (red) and CD19 negative populations (blue). Two regions with overlap of hits in *CRISPR-SURF* and *CRISPR-Millipede* are highlighted as pink (region 1) and light blue (region 2). **c.** Top: TF motifs found at sequences with regulatory potential (high betas). Motifs were filtered based on the expression of the cognate transcription factors in NALM6 cells (>25 CPM) and significant genome-wide CRISPR screen scores (>0.5 or <-0.5) for modulation of *CD19* from a past study^12^. Bottom: PhyloP scores for the *CD19* enhancer show that *CRISPR-Millipede* hits are highly conserved. **d.** PyDESEQ2 analysis: Differentially enriched alleles in CD19 negative or high populations in both screens. Pink dots correspond to alleles containing identified *CRISPR-Millipede* hits in region 1 (151A>G in the ABE8e-SpRy screen and 154G>A in the evoCDA screen). Light blue dots correspond to alleles containing identified *CRISPR-Millipede* hits in region 2 (223A>G, 230A>G in the ABE8e-SpRy screen, 227T>C, 229T>C in the evoCDA screen). Representative alleles are labeled. **e.** *CRISPR-Millipede* visualizations from top to bottom: *board* plot highlighting estimated nucleotide level effects on region 2 (chr16:28930891-28930931). The visualization consists of a heatmap showing *CRISPR-Millipede* effect sizes (square color), PIP (square size) and WT nucleotide with circles for the ABE8e-SpRy screen. Top substitutions with high effect size and PIP include 223A>G and 230A>G. *Board* plot for the evoCDA screen. Top substitutions with high effect size and PIP include 227T>C and 229T>C. Track showing the reference sequence for region 2. Track showing recovered effect sizes for both screens as logo track. Tracks showing candidate TF motifs including SPIB, IKZF1, and PAX5. Tracks showing the editing rate of A>G and T>C and the C>T and G>A substitutions in the two screens (Red: CD19 positive, Blue: CD19 negative).

Next, to overcome the limitations of guide count-based methods in elucidating genotype-phenotype relation with nucleotide-level precision, we employed *CRISPR-CLEAR*. We developed a Bayesian linear regression framework called *CRISPR-Millipede* to effectively distinguish functional nucleotides from bystander edits for both ABE8e and evoCDA screens (**Supplementary Fig. 2b**). This model, leveraging the recent “*millipede”* statistical tool^13^, implements Bayesian Variable Selection for high-dimensional regression and employs an efficient Markov Chain Monte Carlo (MCMC) scheme for inference, allowing it to scale to scenarios with a large number of potential regulatory nucleotides and assigns both effect sizes and Posterior Inclusion Probabilities (PIP) scores to each observed substitution, based on their presence and frequency in the observed alleles across sorted populations.

In contrast to the guide RNA readout model, *CRISPR-Millipede* demonstrates superior capabilities in elucidating genotype-phenotype relation with nucleotide-level precision, a significant improvement over the ∼5-10 base pair resolution of traditional genetic screens. While count-based analysis is biased by differences in the editing rate of individual guide RNAs and underestimates the signal from low-editing guide RNAs, *CRISPR-Millipede* captures editing rates, and identifies regulatory potential based on relative allele frequencies (**Fig. 2b**). This awareness of editing efficiency also enables more accurate estimation of effect sizes and allows the method to disambiguate phenotypic effects more precisely than count-based methods. *CRISPR-Millipede* effectively highlighted clustered substitutions with high and reproducible penetrance on CD19 expression at positions 150 and 223, with high editing rate providing power to detect effects across the tiled regions. Hits identified by *CRISPR-Millipede* map at highly conserved sequences including motifs of transcription factors highly expressed in NALM6 cells, including *MYB*, *SPIB*, *IKZF1* and *PAX5* (**Fig. 2c**).

To leverage the endogenous allelic readout we also adapted the DESeq2 framework^14,15^ to study allele-level enrichment and significance (**Fig. 2d**). This adaptation allowed us to quantify the differential abundance of specific edited alleles between the CD19 positive and negative populations, including the additive effect of multiple substitutions *in cis*.

To better explore the rich output of these screens, we developed novel visualization techniques that create nucleotide “boards” displaying both the PIP and effect size (beta coefficients) for each substitution (**Fig. 2e, top**). Furthermore, we plotted the recovered effect sizes as logo tracks around the clustered substitutions, allowing for direct comparison with the graphical representation of transcription factor motif binding preferences based on position weight matrix (PWM) logos. This enables assessment of the potential impact of each proposed substitution on the binding affinity of each factor. For instance, the logo tracks reveal that observed substitutions often disrupt critical nucleotides for transcription factors like IKZF1/SPIB/PAX5, providing insight into the potential mechanism of action for these variants.

To validate these findings, we designed guide RNAs targeting nucleotides with high phenotypic effect according to CRISPR-Millipede (**Fig. 3a**). In our array of individual guide validations, we observed that sg*145,* a guide RNA disrupting a putative MYB motif, downregulated CD19 expression. Similarly, guide RNAs targeting nucleotides at the IKZF1/SPIB/PAX5 motif in position 220 downregulated CD19 expression as validated by flow cytometry (**Fig. 3b, Supplementary Fig. 3a**). To further characterize the effects of a specific guide, we performed a more detailed analysis on sg*218*. For this guide, we sorted CD19 positive and CD19 negative cells after delivery and sequenced the edited alleles in each population. We observed edited alleles enriched in CD19 negative cells, including substitutions at positions 220, 223, 224 and 230 that are directly linked to CD19 expression (**Fig. 3c**).

**Figure 3:**
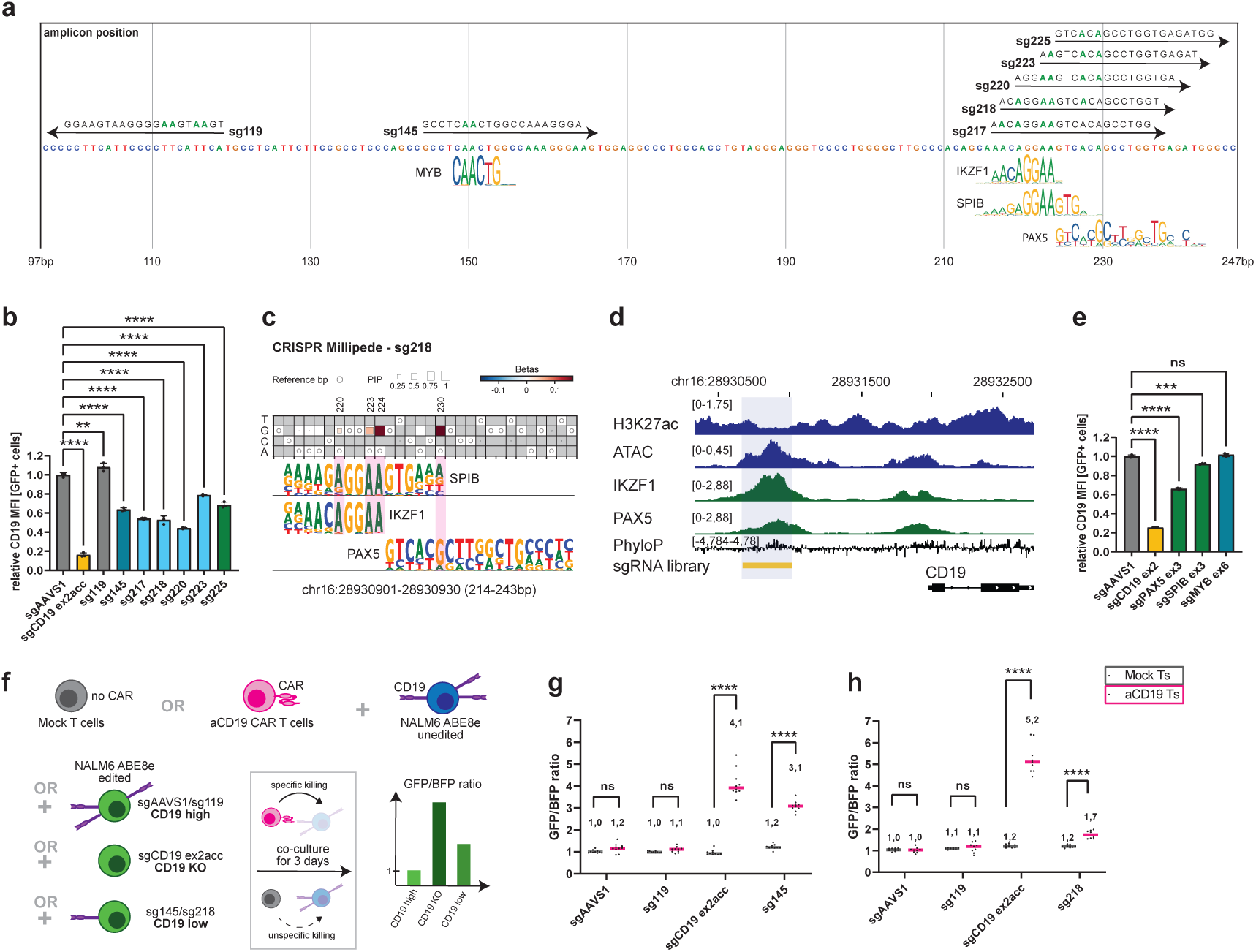
Targeting Millipede hits downregulates CD19 and provides resistance to aCD19-CAR T cells. **a.** Map of the recovered hits by *CRISPR-Millipede* on the *CD19* enhancer sequence, showing TF binding motifs and sgRNAs used for validation experiments. Editable nucleotides are shown in green. **b.** Validation using individual sgRNAs and flow cytometry. While sg119, targeting a neutral sequence within the enhancer showed no effect, editing with sgRNAs 145, 217, 218, 220, 223 and 225 results in downregulation of CD19 MFI compared to sgAAVS1. **c**. Millipede analysis using sg218 highlights nucleotides 220, 223, 224 and 230. **d**. Genomics tracks of the *CD19* locus: the CD19 enhancer is occupied by IKZF1 and PAX5. **e**. Targeting *PAX5* and *SPIB*, but not *MYB*, resulted in downregulation of *CD19*. **f**. Schematics of the CAR-T co-culture experiments. Wild-type, BFP+ NALM6 cells are mixed 1:1 with GFP+ NALM6 cells carrying edits at the *CD19* enhancer and co-cultured with aCD19 CAR T cells or mock T cells. High GFP/BFP ratio indicates that editing facilitates resistance to aCD19-mediated killing. **g.** NALM6 cells edited with sg145 are resistant to aCD19 CAR T. **h**. sg218 provides a milder, yet significant, accumulation of aCD19-resistant cells. One-way ANOVA, replicates are shown in circles (n=3), ** = p<0.01; *** = p<0.001; **** = p<0.0001 (b, e). Multiple unpaired t test, replicates are shown in circles (n=8-10), **** = p<0.000001 (g, h).

To link predicted motifs with their cognate TFs, we targeted the coding sequences of candidate *CD19* regulators *MYB*, *IKZF1*, *SPIB* and *PAX5* using *Sp*Cas9 nuclease and measured CD19 expression. While publicly available ChIP-sequencing data^8^ showed IKZF1 binding at the *CD19* promoter and enhancer (**Fig. 3d**), depletion of IKZF1 did not affect CD19 expression (**Supplementary Fig. 3b**). To confirm these results, we treated *IKZF1*-GFP reporter NALM6 cells with guide RNAs targeting *IKZF1* as well as with Lenalidomide, a potent IKZF1/3 degrader^16^. Loss of GFP indicated effective degradation of IKZF1, while CD19 levels stayed the same, showing that *IKZF1* is not a regulator of *CD19* (**Supplementary Fig. 3c**), despite its binding at the *CD19* enhancer and promoter. Guide RNAs targeting *SPIB*, a lineage-specific TF whose motif is highly similar to that of *IKZF1*, resulted in a 10% decrease in CD19 expression, while targeting *PAX5* showed a pronounced downregulation (∼30%) of CD19. KO of *MYB* did not alter CD19 levels (**Fig. 3e**)

To demonstrate the utility of *CRISPR-CLEAR* in uncovering clinically relevant insights, we investigated the impact of enhancer mutations on CAR-T cell therapy efficacy, a critical issue in cancer treatment. CD19 is a target of CAR-T cell therapy in B-cell malignancies. Despite the high success rate, ∼35% of the B-ALL cases treated with CD19 CAR-T relapses^17^, a proportion of which due to loss or downregulation of the CD19 antigen^18^. To test how substitutions altering the *CD19* enhancer impact physiological expression levels in the context of *CD19* CAR-T, we setup a competition experiment with unedited NALM6 cells (marked with BFP) and genome-edited NALM6 cells (marked with GFP), co-cultured with CD19 CAR-T or mock T-cells, (**Fig. 3f**). After 3 days of co-culture, we observed that targeting the *CD19* exon 2 acceptor splice site resulted in accumulation of cells that were resistant to CAR-T-mediated killing, in line with known variants found at the *CD19* splice site in CAR-T relapses, while sg*119*, targeting a neutral sequence of the *CD19* enhancer did not provide any resistance advantage. Using CAR-T cells derived from independent donors, we observed substantial outgrowth of cells that were resilient to CAR-T after editing with sg*145* and sg*218* (**Fig. 3f-h and Supplementary Fig. d-f**). This suggests that genetic or epigenetic alterations at the *CD19* enhancer might be relevant in the context of CD19-targeted immunotherapies.

In conclusion, our *CRISPR-CLEAR* approach has enabled high-resolution mapping of functional elements within the *CD19* enhancer, revealing specific nucleotides crucial for CD19 expression. We have demonstrated that alterations in these regulatory regions can impact CD19 expression levels and potentially influence the efficacy of CAR-T cell therapies. This study not only provides insights into CD19 regulation but also showcases the power of *CRISPR-CLEAR* in dissecting regulatory elements at single-nucleotide resolution, opening new avenues for understanding gene regulation and improving targeted therapies.

## Code Availability

*CRISPR-Millipede* source code is available at https://github.com/pinellolab/CRISPR-millipede-target. CRISPR-Correct source code is available at https://github.com/pinellolab/CRISPR-Correct. The notebooks and scripts used to generate the figures and analyses presented in the manuscript have been deposited here at https://zenodo.org/doi/10.5281/zenodo.13737736.

## Data Availability

The data used in this manuscript have been provisionally deposited on https://zenodo.org/doi/10.5281/zenodo.13737736.

## Acknowledgements

The authors thank funding from 1R35HG010717–01 (L.P., D.B.), UM1HG012010 (L.P., D.B.), MGH Research Scholar Award 2024-2029 (L.P.). The Seruggia lab is supported by funding from the European Research Council (ERC) under the European Union’s Horizon 2020 research and innovation program (grant agreement 947803), by funding from the Austrian Science Fund (FWF, project P36302) and Marie Sklodowska-Curie (grant agreement 101061151 to A.P.K). We acknowledge Jonathan Hsu and Nafiz Hamid for their initial analyses and discussions that inspired this project. We also thank Manfred Lehner and Elise Sylvander for providing mock and CAR T cells. We thank the CCRI FACS Core Unit for sorting and the CeMM Biomedical Sequencing Facility for sequencing. Finally, we thank members of the Pinello, Seruggia and Bauer lab for providing important feedback on this project.

## Author Contributions

L.P., D.S., and D.B. conceived *CRISPR-CLEAR*. D.S. led the experimental execution and data generation with input from D.B. and L.P. S.W. performed the screens with input from T.W. and M.S., A.K. generated cell lines, S.W. and A.P.K performed characterization of the CD19 enhancer and CAR T experiments.

B.B., Z.M.P., E.C., and J.D. developed the *CRISPR-Millipede* package, with M.J. and L.P. advising on its design and implementation, along with input from D.S. and D.B.

J.D. and Z.M.P. led the *CRISPR-SURF* analyses, and E.C. led the DESeq2 analyses. B.B., Z.M.P., M.J., E.C., and J.D. processed and analyzed the data. L.P., D.S., and D.B. provided guidance and supervised the project. D.S. and L.P. wrote the manuscript with input from all authors. B.B., S.W., and Z.M.P. contributed equally and agreed that either author can be listed first.

## Competing interests

Competing interests L.P. has financial interests in Edilytics, Inc., Excelsior Genomics, and SeQure Dx, Inc. L.P.’s interests were reviewed and are managed by Massachusetts General Hospital and Partners HealthCare in accordance with their conflict of interest policies. The remaining authors declare no competing interests.

## METHODS

### Cell culture and cell lines

NALM6 cells (CVCL_0092) were cultured in RPMI 1640 (Thermo Fisher Scientific) with 10% Fetal Bovine Serum, 100U/mL Penicillin-Streptomycin and 2mM L-Glutamine (referred to as RPMI complete) at 37°C, 5% CO_2_. HEK293T cells were cultured in DMEM (Thermo Fisher Scientific) with 10% Fetal Bovine Serum, 100U/mL Penicillin-Streptomycin and 2mM L-Glutamine. To achieve stable expression of evoCDA and ABE8e-SpRY, 5×10^4 cells were transduced with lentiviruses and selected with 10ug/mL Blasticidin for five days. Expression of base editors was confirmed by Western Blot using a Cas9 antibody (Thermo Fisher Scientific; #MA1-202,dilution 1:1000).

T cells were a gift from Manfred Lehner (CCRI, Vienna). Briefly, healthy donor buffy coats were purchased from the Austrian Red Cross, primary human T cells were purified (RosetteSep Human T cell Enrichment kit; STEMCELL Technologies), and cryopreserved until further use. For generating CAR-expressing T cells, primary human T cells were thawed and activated (Dynabeads Human T-Activator αCD3/αCD28 beads; Thermo Scientific), followed by transduction with purified lentiviruses (Lenti-X Concentrator; Takara) and puromycin selection. T cells were cultured at a density of 0.3 – 2×10^6^ cells/mL in AIMV medium (Thermo Scientific) supplemented with 2% Octaplas (Blutspendezentrale Wien), 2.5% Hepes (PAN Biotech), 1% glutamine (Gibco) and 200 U/mL recombinant human IL-2 (Peprotech).

### Flow cytometry and cell sorting

CD19 expression was measured using a conjugated antibody (APC Clone SJ25C1; BD Biosciences; #340722). Briefly, 100,000-200,000 cells were incubated with a 1:25 antibody dilution in DPBS for 25 minutes at 4°C. APC Mouse IgG1 κ Isotype Control (BD Biosciences; #554681) was used as control. After staining, cells were washed, resuspended in DPBS and analyzed using a BD LSRFortessa™ Cell Analyzer.

For sorting, 20-50×10^6^ cells were washed in DPBS and stained in 50-300 µl staining mix. After staining, cells were washed, resuspended in DPBS and strained through a 35 µm nylon mesh (Fisher Scientific). FACS was performed with BD FACSAria™ Cell Sorter system and sorted cells were collected in 0.5-1mL DPBS.

### Cloning of guide RNAs and tiling library

Guide RNAs were cloned by Golden Gate Cloning in pLKO5.sgRNA.EFS.GFP plasmid (Addgene #57822) and verified by Sanger Sequencing. Oligonucleotides are listed in **Table 1**. The coding sequence of evoCDA-be4b (gift of David Liu) was cloned by Gibson Assembly into XbaI-BamHI digested LentiCas9-Blast (Addgene #52962). The sequence of ABE8e-SpRY (gift of Ben Kleinstiver) was cloned in a lentivector under the SFFV promoter.

All possible 20-mer sequences tiling the top and bottom strand of the *CD19* enhancer were ordered as oligonucleotide pool (IDT), PCR-amplified and cloned by Gibson Assembly in Esp3I-digested pLKO5.sgRNA.EFS.GFP (Addgene #57822). Library sgRNAs and oligonucleotides are listed in **Table 1**. Quality and completeness of the sgRNA library was assessed by NGS.

### Lentivirus production

HEK293T cells were seeded into 15 cm dishes ∼24 hours prior to transfection. Cells were transfected at 80% confluency in 16mL of media with 8.75μg of VSVG, 16.25μg of psPAX2, and 25μg of the lentiviral vector, using 150μg of linear polyethylenimine (PEI) (Sigma-Aldrich). Media was changed to fresh media 16–24 hours post-transfection. Lentiviral supernatant was collected at 48 and 72 hours post-transfection and subsequently concentrated by ultracentrifugation (24000 rpm, 4°C, 2 hours) in a 20% sucrose gradient.

### Base editor screen in NALM6 cells

For base editor screens, 5–7.5×10^6^ NALM6 cells with stable expression of evoCDA or ABE8e-SpRY base editors were transduced at 0.3 MOI to ensure single viral integrations per cell. A titration experiment was performed to determine the amount of virus required to achieve a transduction rate of 30%. A 1000x representation was maintained throughout the screening. Seven days post transduction cells were collected, counted, and processed for FACS sorting. Cells with lentiviral library integrants were selected by gating for GFP. CD19 negative and CD19 positive cells were sorted and genomic DNA was isolated with DNeasy Blood & Tissue Kit (Qiagen). Guide RNAs were enumerated by NGS as previously reported^19^. Distributions of edits within a 346 bp fragment of the *CD19* enhancer were obtained by targeted amplicon sequencing using MiSeq at PE250 and PE300 configuration. Screenings were performed in biological triplicates.

### Quantification of CD19 molecules per cell

For the quantification of CD19 molecules per cell, the BD Quantibrite™ Beads PE Fluorescence Quantitation Kit (BD Biosciences; #340495) was used and CD19 expression in cells was assessed using a PE-conjugated antibody (PE anti-human CD19; clone HIB19; Biolegend; #302208). Briefly, 100,000-200,000 cells were washed in FACS buffer, incubated in blocking solution (10% human serum (Sigma; H4522-20ML) in FACS buffer) for 10 minutes at 4°C, and stained with a 1:100 antibody dilution for 25 minutes at 4°C. PE Mouse IgG1 κ Isotype Control (clone MOPC-21; Biolegend; #400111) was used as control. After two more rounds of washing in the FACS buffer, cells were analyzed using a BD LSRFortessa™ Cell Analyzer. Molecule per cell quantifications were done via linear regression according to the manufacturer’s instructions.

### NALM6-T cell co-culture experiments

NALM6 ABE8e cells were seeded at 500,000 cells per mL in 24 well plates, transduced with concentrated lentiviruses expressing BFP alone or GFP in combination with a sgRNA, and sorted based on BFP or GFP expression 72h post transduction. For co-culture experiments, 25k BFP-and 25k GFP-positive cells were mixed with either mock (non-transduced) or aCD19-expressing CAR T cells in an effector-target-ratio of 0.2:1 and cultured in RPMI media containing 200 U/mL recombinant human IL-2, in U-shaped 96 well plates for 3 days. For flow cytometry, cells were washed in DPBS containing 1% Albunorm (Octapharma; 200g/L) and 0.2% NaN_3_ from a 10% solution and measured using a BD LSRFortessa™ Cell Analyzer.

### Sequencing and demultiplexing

Amplicon-sequencing of endogenous allele samples was conducted on Illumina MiSeq using paired-end 250 and 300 bp and 30% PhiX. Initial demultiplexing for these samples was carried out by the Biomedical Sequencing Facility (BSF) at the Research Center for Molecular Medicine (CeMM). The count samples were sequenced on a NovaSeq SP system, utilizing half of a flow cell with paired-end 100 bp reads. The sequencing core facility performed the primary demultiplexing based on the P7 index. Subsequently, our laboratory conducted a secondary demultiplexing step using the P5 index, which is located internally within the read.

### Guide RNA Mapping

For each demultiplexed FASTQ, “raw” guide mapping was performed using CRISPR-SURF Count (docker version pinellolab/crisprsurf:crispr_clear_v1) in base-editing mode. To address possible self-editing of the guide RNA cassette (especially in ABE8e-SpRY samples), we developed *CRISPR-Correct* (version 0.0.41) which maps the observed guide RNA sequence to an inferred guide RNA sequence that possesses the lowest sequence hamming distance, therefore allowing for possible mismatches between the observed and mapped sequence. Both “raw” and “self-editing corrected” counts were utilized for downstream analysis.

### CRISPR-SURF analysis of guide RNA counts

To identify significant regions based on the mapped guide RNA counts, we ran CRISPR-SURF Deconvolution on both raw and self-editing corrected ABE8e and evoCDA datasets. CRISPR-SURF Deconvolution was run in base-editing mode with a lambda value of 1. For the evoCDA data, guide RNAs with an –NGG PAM and a cytosine within the expected editing window (positions 0 to 14 of the protospacer) were designated as observation guide RNAs, while guide RNAs not meeting these criteria were designated as negative control guide RNAs. The perturbation range for evoCDA was set to 13 based on available data on evoCDA editing activity across six HEK293T sites^9^. For the ABE8e data, guide RNAs that lack an adenine in the expected editing window (positions 2 to 9) were designated as negative controls, and the perturbation range was set to 6, consistent with the known perturbation window for ABE8e^20^.

### CRISPR-Millipede analyses of endogenous alleles

The *CRISPR-Millipede* pipeline consists of five steps taking the demultiplexed FASTQs from amplicon-sequencing of the endogenous alleles as input. The details on the specific steps are provided in the following subsection (**Supplementary Fig. 2b**).

#### 1. CRISPResso2

After targeted amplicon-sequencing of the endogenous alleles, the paired-end FASTQs were quality controlled with FastQC^21^ version 0.12.0 and MultiQC^22^ version 1.15 to ensure sufficient sequencing quality of our samples. Next, the paired-end FASTQs were passed into CRISPResso2 version 2.1.3 for read merging, alignment, and quantification of the edited alleles by running the following command: CRISPRessoBatch –bs {FASTQ_FILENAME} –a {AMPLICON_SEQUENCE} –an cd19 –q 30 –-exclude_bp_from_left 3 –-exclude_bp_from_right 3 –-no_rerun-n {SCREEN_NAME} –-min_frequency_alleles_around_cut_to_plot 0.001 –-max_rows_alleles_around_cut_to_plot 500 –p 20 –-plot_window_size 4 –-base_editor_output –w 0 –bo {OUTPUT_DIRECTORY}

#### 2. Processing and encoding of alleles

To identify functional variants from the CRISPResso2 outputs, we developed a Python package called *CRISPR-Millipede* (version 0.0.89) to process the CRISPResso2 allele tables and perform statistical modeling. Specifically, the CRISPResso2 allele frequency tables were encoded into a feature indicator matrix containing columns for all possible REF>ALT variants across the amplicon and rows for all unique alleles. The first and last 20 positions of the amplicon were removed from the encoding matrix due to sequencing error background. Only canonical REF>ALT variants were included in the encoding matrix depending on the editor used (A>G and T>C for ABE8e samples, C>T and G>A for evoCDA samples). Since evoCDA-NGG is only expected to edit for protospacers with an –NGG PAM, only variants within the canonical editing window (protospacer position 6±7) of the –NGG PAM guide RNAs were included in the encoding matrix.

#### 3. Per-variant modeling of encoded alleles

The matrices are then modeled by millipede^13^ version 0.1.2 (PyPi package millipede-regression), a PyTorch-based Bayesian variable selection model, to generate beta coefficients and posterior inclusion probabilities for each feature. Encoded alleles that do not possess at least two replicates with a read count greater than 2 in at least one population (presort, *CD19* positive, or *CD19* negative) were filtered out to reduce dataset noise. The model was trained using the NormalLikelihoodVariableSelector with the response set to 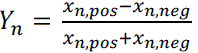 where *x_n,pos_* and *x_n,neg_* are the normalized read count in the *CD19* positive and *CD19* negative populations for each encoded allele *n*, respectively. Normalisation is performed by dividing the raw read count by the total read count for each sample. To account for the higher variance in the response variable caused by alleles with lower read counts, we introduced a scaling factor, called *sigma scale*, which adjusts for the increased uncertainty associated with lower read counts. The sigma scale factor is 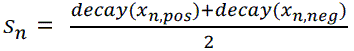 where 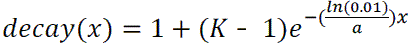, therefore each encoded allele *n* will possess a different relative standard deviation depending on *x_n,pos_*, *x_n,neg_*, *a (*representing the normalized read count where the response noise stabilizes), and *K (*representing the level of response noise for low read counts vs. *a* read counts). For the ABE8e and evoCDA screen samples, K is set to 5 and a is set to 0.0005. For the ABE8e sg218-only sample, K is set to 10 and a is set to 0.0001. This adjustment was made because the score variance for lower read counts in the sg218-only sample is approximately 10 times higher compared to the point where the variance stabilizes. Replicates are modeled jointly by including multiple intercept variables indicating which replicate an encoded allele/response pair belongs to. Millipede attempts to specify a posterior model 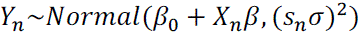 where the β vector contains the set of included features after Bayesian variable selection, thereby producing beta coefficients and posterior inclusion probabilities for each feature.

#### 4. DESeq2 analysis of endogenous alleles

To calculate the differential distribution of each allele in the *CD19* positive and *CD19* negative populations, the encoding matrix from *CRISPR-Millipede* along with a design matrix describing the conditions of each sample (replicate and sorting population) were passed into pyDESeq2^14,15^ version 0.4.10, a statistical package used to calculate differential expression of alleles between the *CD19* positive and *CD19* negative populations. Cook’s distance was calculated and outliers were refit to the model. In addition, log fold change shrinkage was applied and p-values were adjusted for multiple testing. The log fold change and adjusted p-value were plotted for the alleles observed across the cellular populations.

#### 5. Plotting

*CRISPR-SURF* and CRISPR-Millipede plots were generated using jdgenometracks version 0.1.60 (PyPi package jdgenometracks), available at https://github.com/justin-delano/jdgenometracks. Board plots were generated using a custom script.

### Motif analysis

All motif PWMs were obtained from the JASPAR database^23^ (v24). Motifs were scanned using MOODS^24^.

## Supplementary Notes

**Supplementary Figure 1.**
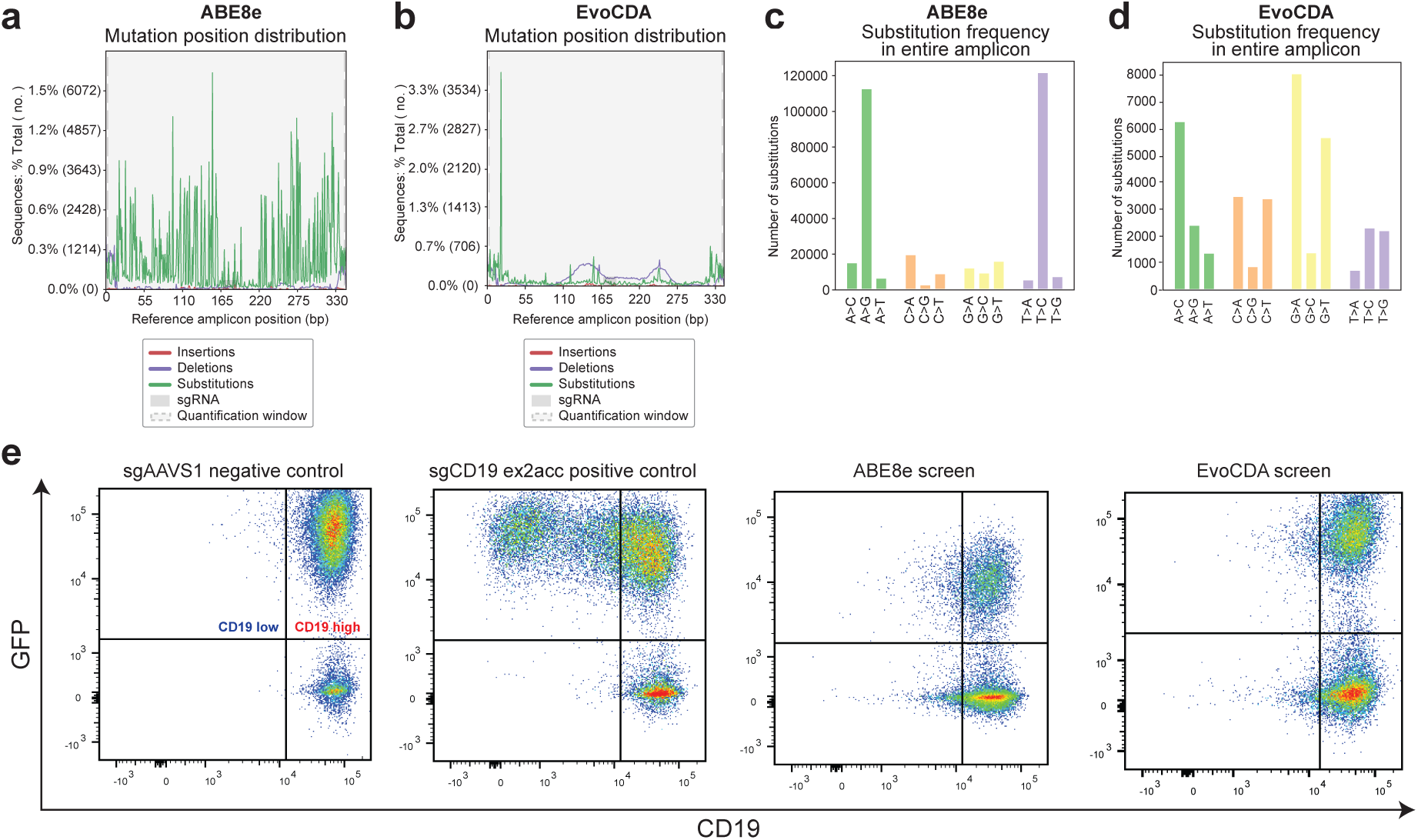
**a**. Distribution and nature of edits at the *CD19* enhancer in NALM6-ABE8e-SpRY cells transduced with the tiling library. ABE8e-SpRY installs single nucleotide substitutions at high density **b**. Editing with evoCDA is sparse and results in substitutions and *indels*. **c**. ABE8e-SpRY introduces specifically A-T>G-C edits. **d.** evoCDA introduces multiple types of substitutions. **e**. Gating strategies used for the FACS sorting. Transduced, GFP positive cells are gated and sorted in CD19 positive and low. As positive control, a guide targeting the adenine of CD19 exon 2 splice acceptor site was used.

**Supplementary Figure 2.**
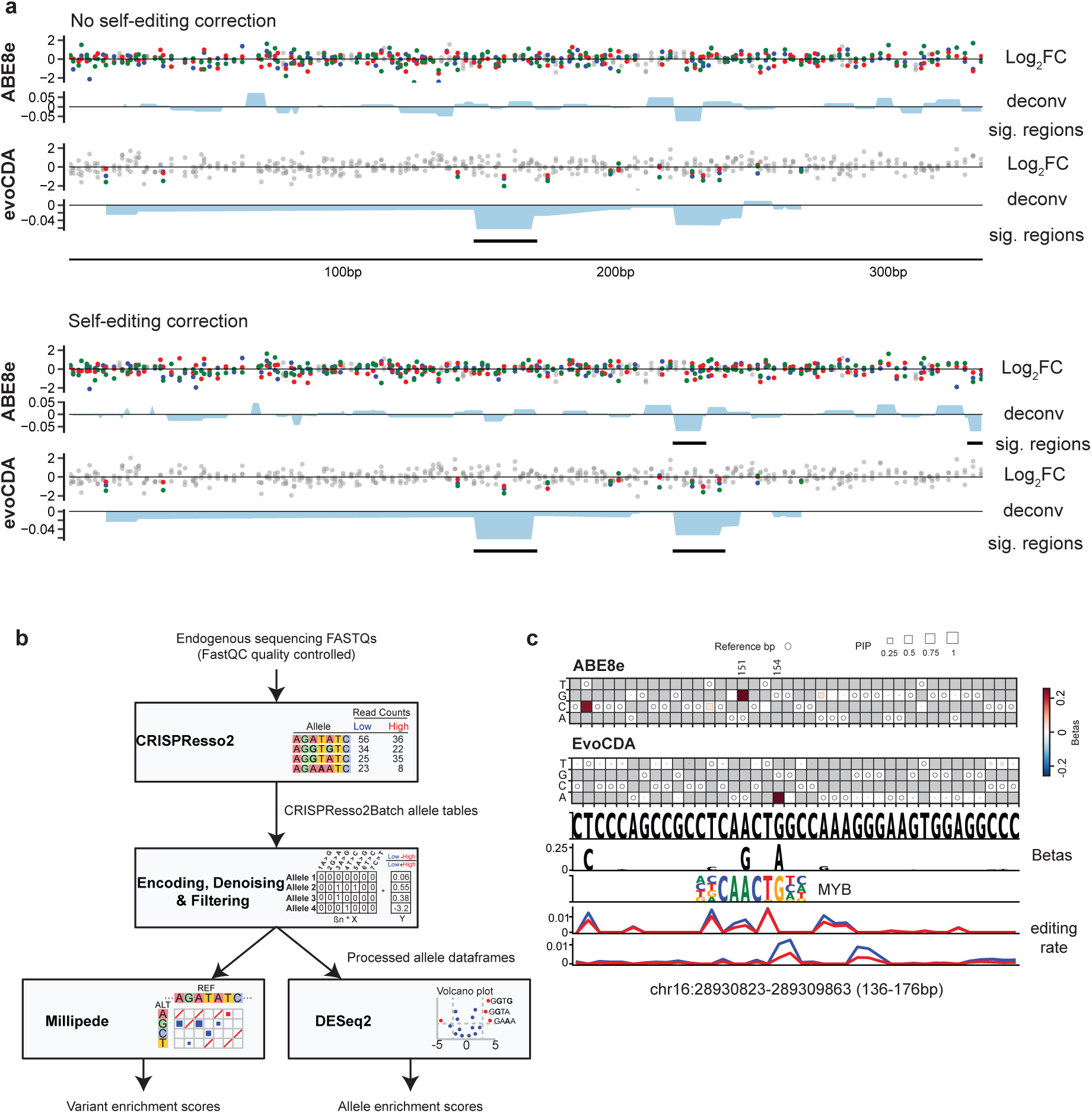
**a**. Comparison of *CRISPR-SURF* analysis from raw vs. self-editing corrected counts. **b**. Schematic of *CRISPR-Millipede* workflow. **c**. *CRISPR-Millipede* plot highlighting region 1 (chr16:28930823-28930863). Heatmap showing *CRISPR-Millipede* effect sizes (square color) and PIP (square size) for the ABE8e-SpRY screen. Top variants with high effect size and PIP include 151A>G. Heatmap showing *CRISPR-Millipede* beta-coefficients (square color) and PIP (square size) for the evoCDA screen. Top variants with high effect size and PIP include 154G>A. Track showing the reference sequence for region 1. Recovered effect sizes are shown as logo tracks. Track showing MYB as candidate TF motif and the editing rate of A>G and T>C and the C>T and G>A substitutions in the two screens (Red: CD19 positive, Blue: CD19 negative).

**Supplementary Figure 3.**
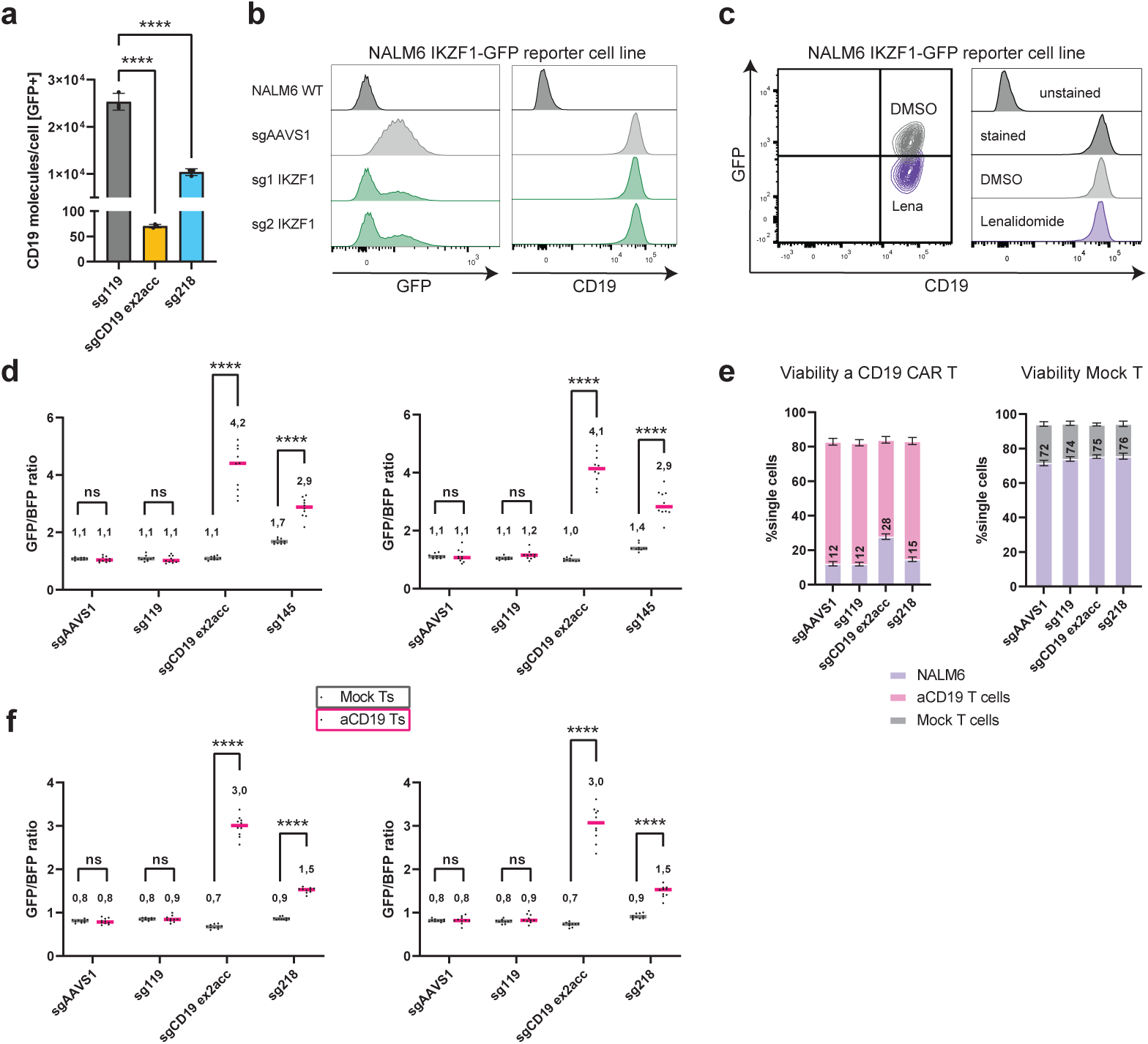
**a**. Quantification of CD19 molecules upon targeting *AAVS1*, *CD19* exon 2 acceptor splice site and sg*218* at the CD19 enhancer. **b.** KO of *IKZF1* in NALM6 IKZF1-GFP cells show depletion of GFP (left) but not in CD19 levels (right). **c**. NALM6 IKZF1-GFP treated with Lenalidomide shows loss of GFP without changes in CD19 expression by flow cytometry. **d,f**. aCD19 CAR competition assay using two additional T cell donors. **e**. Viability plots of NALM6 cells co-cultured with aCD19 CAR or mock T cells. One-way ANOVA, replicates are shown as circles (n=3), **** = p<0.0001 (a). Multiple unpaired t test, replicates are shown as circles (n=10), **** = p<0.000001 (d,f).

## References

1. Nasser J, Bergman DT, Fulco CP, Guckelberger P, Doughty BR, Patwardhan TA, Jones TR, Nguyen TH, Ulirsch JC, Lekschas F, Mualim K, Natri HM, Weeks EM, Munson G, Kane M, Kang HY, Cui A, Ray JP, Eisenhaure TM, Collins RL, Dey K, Pfister H, Price AL, Epstein CB, Kundaje A, Xavier RJ, Daly MJ, Huang H, Finucane HK, Hacohen N, Lander ES, Engreitz JM. Genome-wide enhancer maps link risk variants to disease genes. Nature. 2021 May;593(7858):238–243. PMCID: PMC9153265

2. Canver MC, Smith EC, Sher F, Pinello L, Sanjana NE, Shalem O, Chen DD, Schupp PG, Vinjamur DS, Garcia SP, Luc S, Kurita R, Nakamura Y, Fujiwara Y, Maeda T, Yuan GC, Zhang F, Orkin SH, Bauer DE. BCL11A enhancer dissection by Cas9-mediated in situ saturating mutagenesis. Nature. 2015 Nov 12;527(7577):192–197. PMCID: PMC4644101

3. Gasperini M, Hill AJ, McFaline-Figueroa JL, Martin B, Kim S, Zhang MD, Jackson D, Leith A, Schreiber J, Noble WS, Trapnell C, Ahituv N, Shendure J. A genome-wide framework for mapping gene regulation via cellular genetic screens. Cell. Elsevier BV; 2019 Jan 10;176(1-2):377–390.e19. PMCID: PMC6690346

4. Clement K, Hsu JY, Canver MC, Joung JK, Pinello L. Technologies and Computational Analysis Strategies for CRISPR Applications. Mol Cell. 2020 Jul 2;79(1):11–29. PMCID: PMC7497852

5. Fulco CP, Nasser J, Jones TR, Munson G, Bergman DT, Subramanian V, Grossman SR, Anyoha R, Doughty BR, Patwardhan TA, Nguyen TH, Kane M, Perez EM, Durand NC, Lareau CA, Stamenova EK, Aiden EL, Lander ES, Engreitz JM. Activity-by-contact model of enhancer-promoter regulation from thousands of CRISPR perturbations. Nat Genet. 2019 Dec;51(12):1664–1669. PMCID: PMC6886585

6. IGVF Consortium. Deciphering the impact of genomic variation on function. Nature. 2024 Sep;633(8028):47–57. PMID: 39232149

7. Yao D, Tycko J, Oh JW, Bounds LR, Gosai SJ, Lataniotis L, Mackay-Smith A, Doughty BR, Gabdank I, Schmidt H, Youngworth I, Andreeva K, Ren X, Barrera A, Luo Y, Siklenka K, Yardımcı GG, The ENCODE4 Consortium, Tewhey R, Kundaje A, Greenleaf WJ, Sabeti PC, Leslie C, Pritykin Y, Moore JE, Beer MA, Gersbach CA, Reddy TE, Shen Y, Engreitz JM, Bassik MC, Reilly SK. Multi-center integrated analysis of non-coding CRISPR screens [Internet]. bioRxiv. 2022 [cited 2024 Sep 7]. p. 2022.12.21.520137. Available from: https://www.biorxiv.org/content/10.1101/2022.12.21.520137v1

8. Okuyama K, Strid T, Kuruvilla J, Somasundaram R, Cristobal S, Smith E, Prasad M, Fioretos T, Lilljebjörn H, Soneji S, Lang S, Ungerbäck J, Sigvardsson M. PAX5 is part of a functional transcription factor network targeted in lymphoid leukemia. PLoS Genet. 2019 Aug;15(8):e1008280. PMCID: PMC6695195

9. Thuronyi BW, Koblan LW, Levy JM, Yeh WH, Zheng C, Newby GA, Wilson C, Bhaumik M, Shubina-Oleinik O, Holt JR, Liu DR. Continuous evolution of base editors with expanded target compatibility and improved activity. Nat Biotechnol. 2019 Sep;37(9):1070–1079. PMCID: PMC6728210

10. Walton RT, Christie KA, Whittaker MN, Kleinstiver BP. Unconstrained genome targeting with near-PAMless engineered CRISPR-Cas9 variants. Science. 2020 Apr 17;368(6488):290–296. PMCID: PMC7297043

11. Hsu JY, Fulco CP, Cole MA, Canver MC, Pellin D, Sher F, Farouni R, Clement K, Guo JA, Biasco L, Orkin SH, Engreitz JM, Lander ES, Joung JK, Bauer DE, Pinello L. CRISPR-SURF: discovering regulatory elements by deconvolution of CRISPR tiling screen data. Nat Methods. 2018 Dec;15(12):992–993. PMCID: PMC6620603

12. Witkowski MT, Lee S, Wang E, Lee AK, Talbot A, Ma C, Tsopoulidis N, Brumbaugh J, Zhao Y, Roberts KG, Hogg SJ, Nomikou S, Ghebrechristos YE, Thandapani P, Mullighan CG, Hochedlinger K, Chen W, Abdel-Wahab O, Eyquem J, Aifantis I. NUDT21 limits CD19 levels through alternative mRNA polyadenylation in B cell acute lymphoblastic leukemia. Nat Immunol. 2022 Oct;23(10):1424–1432. PMCID: PMC9611506

13. Jankowiak, M. Bayesian Variable Selection in a Million Dimensions. In: Ruiz F, Dy J, van de Meent JW, editors. Proceedings of The 26th International Conference on Artificial Intelligence and Statistics. PMLR; 25--27 Apr 2023. p. 253–282.

14. Love MI, Huber W, Anders S. Moderated estimation of fold change and dispersion for RNA-seq data with DESeq2. Genome Biol. 2014;15(12):550. PMCID: PMC4302049

15. Muzellec B, Teleńczuk M, Cabeli V, Andreux M. PyDESeq2: a python package for bulk RNA-seq differential expression analysis. Bioinformatics [Internet]. 2023 Sep 2;39(9). Available from: 10.1093/bioinformatics/btad547 PMCID: PMC10502239

16. Krönke J, Udeshi ND, Narla A, Grauman P, Hurst SN, McConkey M, Svinkina T, Heckl D, Comer E, Li X, Ciarlo C, Hartman E, Munshi N, Schenone M, Schreiber SL, Carr SA, Ebert BL. Lenalidomide causes selective degradation of IKZF1 and IKZF3 in multiple myeloma cells. Science. 2014 Jan 17;343(6168):301–305. PMCID: PMC4077049

17. Maude SL, Laetsch TW, Buechner J, Rives S, Boyer M, Bittencourt H, Bader P, Verneris MR, Stefanski HE, Myers GD, Qayed M, De Moerloose B, Hiramatsu H, Schlis K, Davis KL, Martin PL, Nemecek ER, Yanik GA, Peters C, Baruchel A, Boissel N, Mechinaud F, Balduzzi A, Krueger J, June CH, Levine BL, Wood P, Taran T, Leung M, Mueller KT, Zhang Y, Sen K, Lebwohl D, Pulsipher MA, Grupp SA. Tisagenlecleucel in Children and Young Adults with B-Cell Lymphoblastic Leukemia. N Engl J Med. 2018 Feb 1;378(5):439–448. PMCID: PMC5996391

18. Orlando EJ, Han X, Tribouley C, Wood PA, Leary RJ, Riester M, Levine JE, Qayed M, Grupp SA, Boyer M, De Moerloose B, Nemecek ER, Bittencourt H, Hiramatsu H, Buechner J, Davies SM, Verneris MR, Nguyen K, Brogdon JL, Bitter H, Morrissey M, Pierog P, Pantano S, Engelman JA, Winckler W. Genetic mechanisms of target antigen loss in CAR19 therapy of acute lymphoblastic leukemia. Nat Med. 2018 Oct;24(10):1504–1506. PMID: 30275569

19. Seruggia D, Oti M, Tripathi P, Canver MC, LeBlanc L, Di Giammartino DC, Bullen MJ, Nefzger CM, Sun YBY, Farouni R, Polo JM, Pinello L, Apostolou E, Kim J, Orkin SH, Das PP. TAF5L and TAF6L Maintain Self-Renewal of Embryonic Stem Cells via the MYC Regulatory Network. Mol Cell. 2019 Jun 20;74(6):1148– 1163.e7. PMCID: PMC6671628

20. Arbab M, Shen MW, Mok B, Wilson C, Matuszek Ż, Cassa CA, Liu DR. Determinants of Base Editing Outcomes from Target Library Analysis and Machine Learning. Cell. 2020 Jul 23;182(2):463–480.e30. PMCID: PMC7384975

21. Babraham bioinformatics – FastQC A quality control tool for high throughput sequence data [Internet]. [cited 2024 Sep 8]. Available from: http://www.bioinformatics.babraham.ac.uk/projects/fastqc/

22. Ewels P, Magnusson M, Lundin S, Käller M. MultiQC: summarize analysis results for multiple tools and samples in a single report. Bioinformatics. 2016 Oct 1;32(19):3047–3048. PMCID: PMC5039924

23. Rauluseviciute I, Riudavets-Puig R, Blanc-Mathieu R, Castro-Mondragon JA, Ferenc K, Kumar V, Lemma RB, Lucas J, Chèneby J, Baranasic D, Khan A, Fornes O, Gundersen S, Johansen M, Hovig E, Lenhard B, Sandelin A, Wasserman WW, Parcy F, Mathelier A. JASPAR 2024: 20th anniversary of the open-access database of transcription factor binding profiles. Nucleic Acids Res. 2024 Jan 5;52(D1):D174–D182. PMCID: PMC10767809

24. Korhonen J, Martinmäki P, Pizzi C, Rastas P, Ukkonen E. MOODS: fast search for position weight matrix matches in DNA sequences. Bioinformatics. 2009 Dec 1;25(23):3181–3182. PMCID: PMC2778336

